# Genetic variation in *BnGRP1* contributes to low phosphorus tolerance in *Brassica napus*

**DOI:** 10.1101/2022.07.14.500146

**Authors:** Ping Xu, Haiyuan Li, Ke Xu, Xiaoyu Cui, Zhenning Liu, Xiaohua Wang

**Author notes:** **Corresponding author:** Dr. Xiaohua Wang. **Email address: Ping Xu**. **Haiyuan Li**. **Ke Xu**. **Xiaoyu Cui**. **Zhenning Liu**. **Xiaohua Wang**.

## Abstract

The lack of phosphorus (P) is a major environmental factor affecting rapeseed root growth and development. For breeding purposes, it is crucial to identify the molecular mechanisms of root system architecture (RSA) traits underlying low P tolerance in rapeseed. Using genome-wide association study (GWAS), transcriptome and re-sequencing analyses were done on 400 rapeseed cultivars, the natural variations of glycine-rich protein gene, *BnGRP1*, in response to low P tolerance. Based on 11 SNP mutations in the *BnGRP1* sequence, ten types of haplotypes (Hap) were formed. Compared with the other types, the cultivar of the *BnGRP1*^*Hap1*^ type in the panel demonstrated the longest root length and heaviest root weight. Over-expression of *BnGRP1*^*Hap1*^ in rapeseed depicted the ability to enhance the resistance of rapeseed in its response to low P tolerance. CRISPR/Cas9-derived *BnGRP1*^*Hap1*^ knockout mutations in rapeseed could lead to sensitivity to low P stress. Furthermore, *BnGRP1*^*Hap1*^ influenced the expression of phosphate transporter 1 (*PHT1*) genes associated with P absorption. Overall, the findings of this study highlight the new mechanisms of *GRP1* genes in enhancing the low P tolerance in rapeseed.

## Introduction

Due to its widespread use in edible and industrial oils, canola-type rapeseed (*Brassica napus*) is one of the most important oil crops globally (Tan et al., 2022). In rapeseed, phosphorus (P) plays an important role during the growth period and in sensitivity to low P environments (Wang et al., 2017; Xu et al., 2022). However, most P in soil is fixated by iron, aluminum, or calcium in the soil (Yamaji et al., 2017; Poirier et al., 2022), preventing it from being absorbed and utilized by rapeseed. Because of the limitations in rock phosphate for the production of P fertilizers, breeding high P-efficiency rapeseed varieties and revealing the molecular mechanism are key strategies in reducing P fertilizers (Wang et al., 2017. As the most important organ for nutrient absorption, root system architecture (RSA) traits are important to achieve genetic improvements in rapeseed (Xu et al., 2022). As a result of low P tolerance, plants have evolved a series of adaptive mechanisms to improve P acquisition efficiency, such as increased root surfer area, increased primary root length (PRL), and increased root and shoot dry weight (Wissuwa, 2005; Lynch, 2011; Mori et al., 2016; Zhang et al., 2016). Therefore, identifying the genetic loci related to RSA traits and revealing the P-efficient mechanism is very important in rapeseed breeding.

Low P tolerance traits are quantitative traits controlled by multiple genes. In response to the detection of low P tolerance, the traditional strategies for RSA traits include linkage analysis and genome-wide association study (GWAS) (Wang et al., 2017; Xu et al., 2022). Several RSA trait loci (QTLs) have been detected in various plants. Not only that, many major QTL/QTLs related to RSA traits, P uptake, and P efficient traits in *A. thaliana*, rice, bean, and *B. napus* have been detected in segregating populations constructed by bi-parental crossing. For instance, one QTL-related root trait under P was detected in the *A. thaliana* recombinant inbred lines (RILs) population (Svistoonoff et al., 2007). Next, one QTL related to P uptake was detected in F_3_ near-isogenic lines population (Wissuwa et al., 2002). In addition, two QTLs related to P uptake were identified in F_5:7_-derived RILs (Yan et al., 2004). A total of 64 QTLs related to RSA traits were detected in a doubled haploid population (Zhang et al., 2016). Moreover, hundreds of significant genetic loci related to P efficiency traits and RSA traits were detected in rich genetic variation association panels in soybean (Zhang et al., 2014), rice (Mori et al., 2016), and rapeseed (Wang et al., 2017). The number of significant SNPs/cultivars were 74/192 in soybean, 6/413 in rice, and 285/405 in rapeseed.

Phosphorus uptake in plants is a transport process mediated by high-affinity phosphate transporters (PHT) across the plasma membrane. As PHT is expressed in the cytoplasmic membrane in root tissue of various crops, they could drive P uptake by plants via regulation of the hydrogen ion concentration gradient in the cytoplasmic membrane (DiTusa et al., 2016; Poirier et al., 2022). Therefore, a common mechanism used by plants while adapting to low P tolerance is the up-regulated expression of PHT1 family members (Dai et al., 2022). In the P signaling network, PHT1 family gene expression is regulated by upstream transcription factors and genes (Zhang et al., 2022). Several pathways through which transcription factor phosphate starvation response 1 (PHR1) can regulate *PHT1* expression. First, it directly regulates the expression of the PHT1 gene by binding to the PHR domain of the PHT1 gene (Liang et al., 2014). Secondly, *OsPHR2* can inhibit the ubiquitination of *PHT1* by regulating *OsmiR399* expression and indirectly promote the expression of *PHT1* under low P stress in rice (Chiou and Lin, 2011). Lastly, the *PHT1* gene is regulated specifically in response to low P stress by forming the AtPHR1-AtSPX4 conjugate when P is abundant and releasing *PHR1* when P is deficient in *Arabidopsis* (Lv et al., 2014). In transgenic *Arabidopsis*, over-expression of the zinc-finger transcription factor 6 (*AtZAT6*) and *AtMYB62* was found to inhibit the expression of *AtPHT1;1* and *AtPHT1;4* (Chiou and Lin, 2011). In this study, the glycine-rich protein gene (*BnGRP1*) was overexpressed, and the expression of *BnPHT1;4* and *BnPHT1;7* were increased, but the expression of *BnZAT6* was inhibited in rapeseed.

GRPs are glycine-rich proteins composed of highly repetitive glycine sequences and a signal peptide sequence at the N-terminus (Kim et al., 2008; Wu et al., 2020). Due to their key role in plant stress response and developmental regulation, GRPs have become the topic of a current research hotspot. In a previous study, the over-expression of *BnGRP7* was noted to enhance resistance against cold and drought resistance by regulating cell osmosis through membrane oxidative capacity in *Arabidopsis* (Kim et al., 2008). The homologous gene of *MpGR-RBP1* in *Malus prunifolia* enhanced the resistance of salt stress through influencing on ROS accumulation and stomatal behavior confers salt stress tolerance and protects against oxidative stress in *Arabidopsis* (Tan et al., 2014).. *NtGRP3* promoted root elongation in response to low sulfur stress by regulating the degree of lignification of vascular bundles in tobacco (Znój et al., 2017)..

Although previous studies have detected several significant markers associated with low P tolerance, their molecular mechanisms were unclear. Moreover, knowledge about GRPs response to low P stress was rare at the time of the present study. Therefore, to better understand the genetic control of RSA traits related to low P tolerance, a population of 400 accessions genotyped with *Brassica* 60K SNP arrays and re-sequencing were used for investigating the genetic control of RSA traits enhancement of low P tolerance. The most significant variation related to multiple RSA traits under low P was located in the confidence interval of *BnGRP1*, encoding the glycine-rich protein. A total of 11 SNPs of *the BnGRP1* sequence formed ten haplotypes, and the cultivars with haplotype 1 (*BnGRP1*^*Hap1*^) had the best RSA traits in the association panel. Overexpression and knockout of *BnGRP1*^*Hap1*^ led the transgenic lines towards an enhanced or sensitized response to low P stress, respectively. In addition, *BnGRP1*^*Hap1*^ promoted the expression of *BnPHT1* but inhibited the *BnZAT6* in P signaling network. Overall, the results have increased our understanding of the molecular mechanisms of *GRP1* in enhancing low P tolerance in rapeseed.

## Materials and methods

### Plant materials and phenotype under low P conditions

For GWAS, the natural population of 400 rapeseed accessions provided by Huazhong Agricultural University (Wuhan, China) (Wang et al., 2017) was used. Rapeseed materials P inefficiency cultivar Westar 10 (W10) and P efficient cultivar Zhongshuang 11 (ZS11) were used for transgenics. The transgenic line of *35S::BnGRP1*^*Hap1*^, *Bngrp1*, and *BnGRP1*^*Hap1*^*::GFP* was previously described in Xu et al. (2017).

The cultivars of natural population and transgenic lines were planted in a modified Hogland solution (-S) and phosphorus-free paper culture (-P) at low P (0.005 mmol·L^-1^) conditions during the years 2015, 2016, 2021, and 2022. More than 40 plants were planted following the method described previously by Wang (2017). Certain RSA traits (total root length, TRL; root volume, RV; lateral root number, LRN) of rapeseed were obtained via WinRHIZO root analysis system (http://www.agripheno.com/en/WinRHIZO.html). Other RSA traits (primary root length, PRL; lateral root length, LRL; mean lateral root length, MLRL; lateral root density, LRD) were measured by the ImageJ software (https://imagej.nih.gov/ij/) 30 days after sowing. The root and shoot tissues were separated and dried at 80 °C to a constant weight. The shoot dry weight (SDW) and root dry weight (RDW) were determined. RSA traits and haplotype boxplots were drawn using Origin 8 software (http://www.uone-tech.cn/Origin.html). The probability level and statistical analysis of phenotype were performed by GenStat 18^th^ (https://vsni.co.uk/) and Excel 2017 (http://rjxz.cmgsll.com.cn/html/office/). The difference was significant at 1% probability level (Wang et al., 2017).

### Genotyping by Brassica 60K SNP array and re-sequencing

The DNA of natural population cultivars and transgenic rapeseeds were extracted by a modified hexadecyl trimethyl ammonium bromide (CTAB) method (Xu et al., 2017). Based on the criteria of DNA concentration = 50 ng·μL^-1^, DNA length > 10 Kb, high-quality DNA was utilized for SNP detection through *Brassica* 60K Illumina Infinium SNP array and Illumina NovaSeq 6000 (Illumina Inc, San Diego, USA). The SNPs were mapped to the *Brassica napus* reference cultivar ‘ZS11’ (http://cbi.hzau.edu.cn/bnapus/index.php) using a 50 bp aligning sequence of SNPs and 150 bp reads by local BLASTn (https://blast.ncbi.nlm.nih.gov/Blast.cgi). GWAS and haplotype analysis utilized high-quality SNPs with more than 80% of call frequencies, higher than 5% of minor allele frequencies, and each SNP hitting one physical position.

### GWAS for RSA traits

To correct the spurious association led by genotype-phenotype covariance, the population structure and relative kinship of cultivars in the natural population were used. In this study, the population structure was estimated according to the Bayesian Markov Chain Monte Carlo model (MCMC) in the software of STRUCTURE 2.3.4 (http://web.stanford.edu/group/pritchardlab/home.html). The relative kinship matrix was calculated using SPAGeDi software (https://spagedi.software.informer.com/). The pairwise negative kinship values were set to 0. According to Nei’s genetic distance (Nei, 1972) among cultivars, the Neighbor-Joining tree of the natural population was constructed by MEGA6 software (https://mega6.software.informer.com/). Based on the parameter *r*^*2*^ of pairwise markers, the linkage disequilibrium (LD) decay was evaluated using TASSEL version 5.2.81 (https://www.softpedia.com/get/Science-CAD/TASSEL.shtml).

To detect genetic variations in RSA traits, two models (general linear model, GLM, and mixed linear model, MLM) by GWAS were used. The quantile-quantile (QQ) plot-related expected and observed of each SNP used for appropriate model detection was calculated using TASSEL 5.2.81. The significance analysis of SNPs and traits was based on the threshold of the Bonferroni correction for multiple tests. Based on the –log10(P) of each SNP (P = 1/n, n was the SNP number), the QQ plot and Manhattan plot were drawn by R script (Wang et al., 2017).

### Differential expression genes detection and real-time RT-PCR

Total RNA was extracted from leaf and root tissues of the natural population and transgenic rapeseed using RNAsimple Kit (DP419, TIANGEN, China). The RNA was sequenced using Illumina 2500 platform (Illumina Inc, San Diego, USA). Based on the criteria of a false discovery rate (FDR) less than 1%, *p* values less than 1%, and fold change of NP/LP > 2 or transgenic/WT > 2, the differential expression genes were identified. After reverse transcription, the relative expression of genes was quantified using the 2^-^ΔΔCt method (Xu et al., 2022).

### Haplotype analysis of BnGRP1

The variation in haplotype types could lead to different phenotypes through gene expression or gene sequence mutations. Based on the SNP detected by re-sequencing and comparing sequencing, the haplotypes of *BnGRP1* were analyzed. TASSEL 5.2.81 software was utilized to identify the haplotype blocks access to the natural population. OriginPro 8.0 software was used to draw the boxplots of RSA traits with low P tolerance (Wang et al., 2017).

### Construction of the CRISPR/Cas9 vector and plant transformation

The binary vector system pKSE401 (Chen et al. 2017) with a kanamycin selection marker to target the P efficient *BnGRP1*^*Hap1*^ gene was used. Two target regions were selected to mutate the *BnGRP1*^*Hap1*^ and its homologous copy through the Web-based tool of CRISPR-P 2.0 (http://crispr.hzau.edu.cn/CRISPR2). One was able to target all the copies, and the other site only specifically targeted the copy of *BnGRP1*^*Hap1*^. The primers (G1-DT1-BsF, G1-DT1-F0, G1-DT2-R0, G1-DT2-BsR) used for the construction of the sgRNA vector are listed in the Table S9. The method of hypocotyl transformation mediated by Agrobacterium tumefaciens in *B. napus* (Zhang et al., 2015) was used to transform “ZS11”.

### Sub-cellular location of BnGRP1

The CDS sequence of *BnGRP* ^*Hap1*^ without the termination codon (TAA) was inserted downstream of the double 35S promoter through Xba I - BamH I site in frame with GFP in the pMDC83 vector to generate the *BnGRP1*:GFP fusion constructs and infiltrated into *N. Benthamiana* leaves. All the leaves were collected at 36 h after transformation and the fluorescent signal was detected using confocal microscopy (IX83-FV1200, Olympus).

### GUS staining

Transgenic plants carrying the *BnGRP1* ^*Hap1*^ promoter-GUS fusion construct in the *Arabidopsis* were constructed. Briefly, a 1.8-kb genomic DNA fragment of the 5’ upstream region of *BnGRP1* ^*Hap1*^ amplified by PCR using primers, GRP-4L and GRP-4R (Table S10) were cloned into the pBI121 vector. Transgenic seedlings of 35 d were incubated at 37 °C overnight in 5-bromo-4-chloro-3-indolyl-b-glucuronic acid solution and then cleaned in 75% (v/v) ethanol. The treated seedlings were observed on an Olympus IX-70 microscope equipped with Nomarski Optics (Li et al., 2015).

### The Pi concentration of transgenic rapeseed

Dry shoot/root tissues of rapeseed weights 0.1–0.5 g and 5mL of 1% (v/v) HNO_3_ were digested overnight in the Teflon tank. After 2 h at 80 °C, 2 h at 120 °C, and 4 h at 160 °C, the lid was opened, and the mixture was heated to remove the acid. The digestion solution was washed into a 25 mL volumetric flask, and the volume was made up of 1% (v/v) HNO_3_. The shoot and root tissues were measured using Agilent 710 ICP-OES (Nanjing Webiolotech Biotechnology Co., Ltd, Nanjing, China).

## Results

### Phenotype analysis of RSA traits response to low P stress

Out of 400 natural accessions grown in Hogland’s solution (-S) culture at low P conditions, four RSA traits (root dry weight-RDW, shoot dry weight-SDW, primary root length-PRL, and root shoot ratio-RS ratio) were obtained with three environment repetitions. Under low P stress, 10 RSA traits (mean lateral root length-MLRL, lateral root density-LRD, lateral root length-LRL, lateral root number-LRN, total root length-TRL, root angle, RDW, SDW, PRL, and RS ratio) were collected access phosphorus-free paper culture (-P) (Table 1). All the RSA traits demonstrated wide variation and continuous Gaussian distribution (Fig. 1), indicating that RSA traits were quantitative traits controlled by multiple genes. Furthermore, four traits (RDW, PRL, SDW, and RS ratio) revealed significant positive correlations (*p =* 4.2×10^−7^, R^2^ = 0.4595; *p = 5*.*85*×10^−6^, R^2^ = 0.9687; *p =* 3.43×10^−6^, R^2^ = 0.5929, and *p =* 3.37×10^−5^, R^2^ = 0.1868, respectively) between -S culture and –P culture, under low P stress.

**Fig 1.**
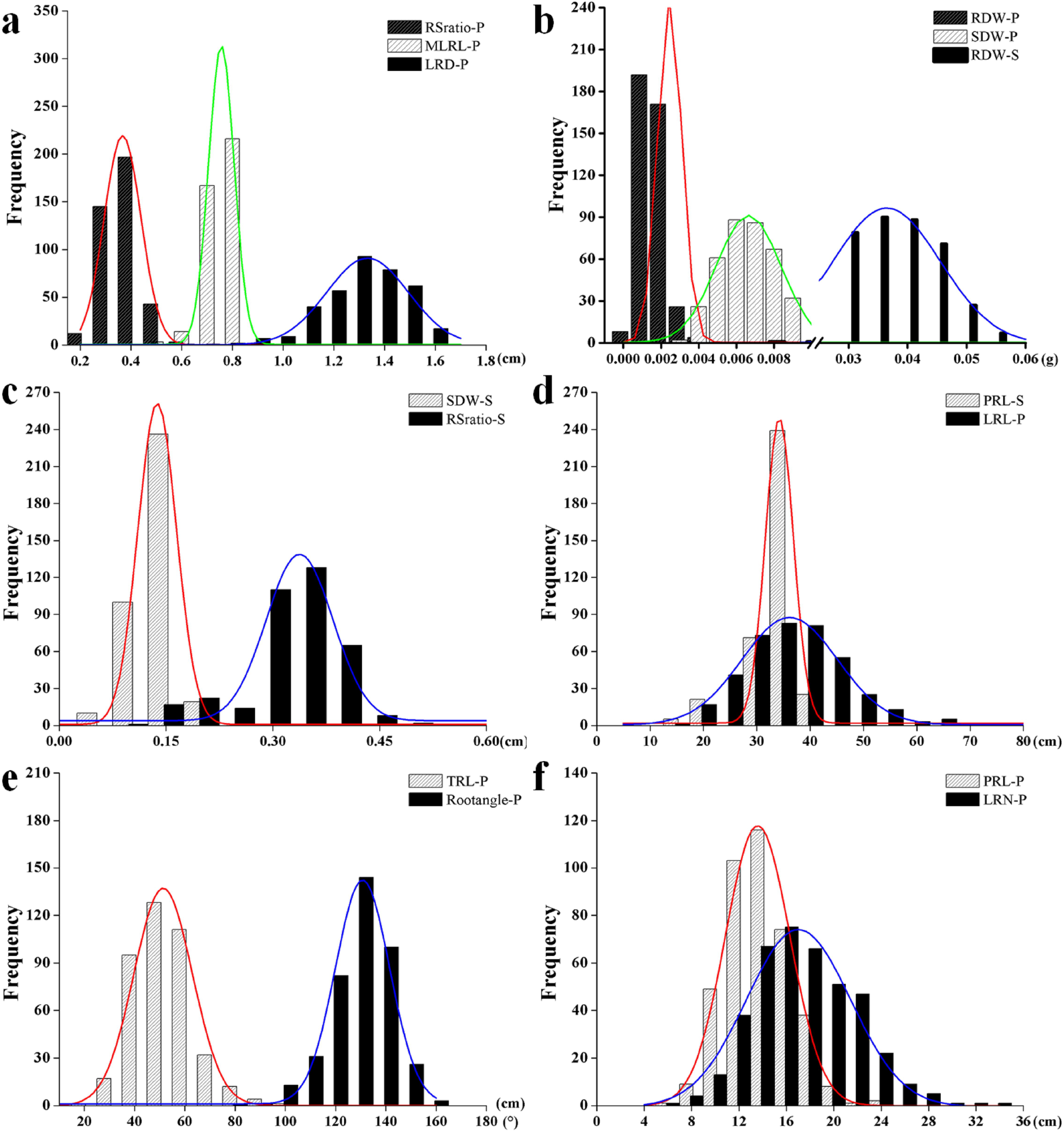

### Genomic variation, population structure, Kinship, and LD decay of rapeseed natural population

In total, 32369 high-quality SNPs were mapped on reference cultivar ZS11. The number of SNPs on 19 chromosomes ranged from 927 on C9 to 2816 on C4, with an average of 1703. The number of SNPs on the A and C sub-genome were 16564 and 15805, respectively (Table S1). The genetic distance is shown in Table S2. The cultivars in the population can be divided into two sub-genomes, P1 with 335 cultivars and P2 with 65 cultivars (Fig.s 2a and b; Table S2). When the relative kinship values were equal to 0, pairwise kinship coefficients were 68%, indicating that the 400 cultivars had a weak relationship in this study (Fig. 2c). Based on the cut-off for squared correlations of allele frequencies (*r*^2^) at 0.1, the LD decay distance of this natural population was 2065 Kb. Moreover, LD decay was significantly different in each chromosome, from 125 Kb on A4 to 8250 Kb on C1 chromosome (Table S3). LD decay of the C genome (8250 Kb) was five times that of A genome (1500 Kb) (Fig. 2d; Table S3).

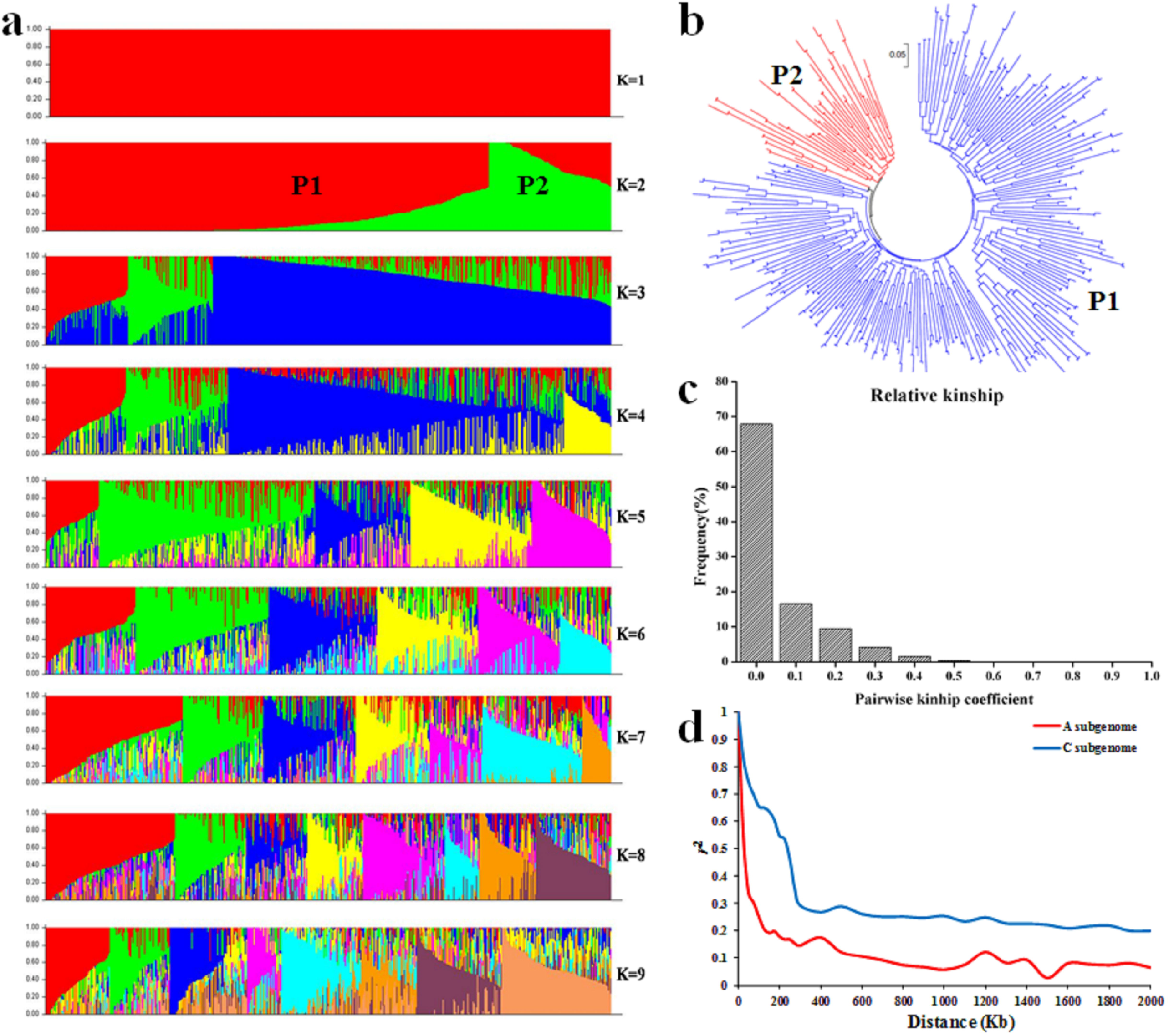

### Genome-wide association mapping of RSA traits in rapeseed

Two common models (GLM and MLM) were used to control the false positive of genotype-phenotype. Based on *p* values < 3.09×10^−5^ (n = 32369) (Table S1), a total of 440 significant SNPs related to ten RSA traits growth at two cultures under low P stress were detected on 19 chromosomes (Fig. 3a; Table S4). QQ plots were used to establish the appropriate model (Table S4). The significant SNP marker (SNP5946) on the A5 chromosome could explain 10.72%, 11.00%, 10.64%, 11.27%, and 11.07% phenotypic variation of LRN, PRL, RDW, SDW, and TRL traits under low P stress (Fig. 3a; Table S4). A total of 1651 SNPs (Table S5) located on A5 were used for a single chromosome association study revealing the key SNPs related to RSA traits in this population. According to *p* < 6.06×10^−4^ (n = 1651), a total of 141 peak SNPs related to RSA traits were identified on the A5 chromosome (Fig. 3b; Table S6). The phenotype variations of LRN, PRL, RDW, SDW, TRL, and LRL under low P stress were explained by the significant SNP (SNP127-A5) at 8.54%, 8.44%, 8.87%, 11.67%, 11.07%, and 10.56%, respectively (Table S6).

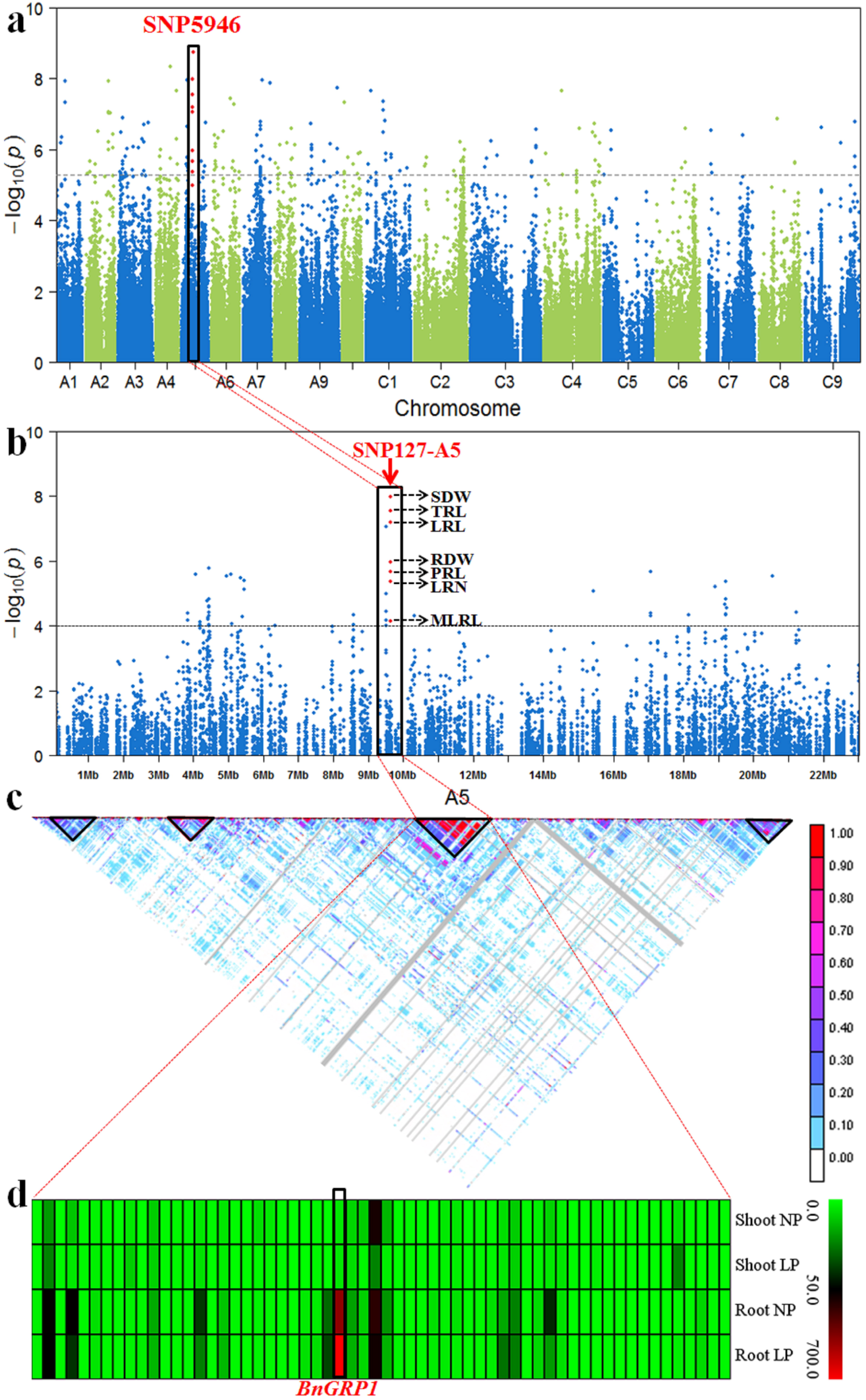

### BnGRP1 was significantly associated with RSA traits related to low P tolerance

Four haplotype blocks were detected on the A5 chromosome, and the significant SNPs (SNP5969 and SNP127-A5) were mapped on the third haplotype block (Fig. 3c). The candidate gene (*BnGRP1*) mapped on the LD decay confidence interval of peak SNP (SNP127-A5) revealed increased expression (from 423.9 to 796.6 FPKM) in root tissue response to low P stress (Fig. 3d; Table S7). Based on the re-sequencing, 27 SNP were detected and mapped on the *BnGRP1* sequence (Fig. 4a; Table S8). Next, 11 of 27 SNPs formed a block with ten haplotype types in the *BnGRP1* gene sequence (Fig.s 4b-d; Table S8). Based on the 11 SNPs in *the BnGRP1* gene region, ten haplotype types were invested in the top five RSA traits (SDW, TRL, LRL, RDW, PRL, LRN, and MLRL) related to *BnGRP1* (Fig. 4e-l). After the relationship between haplotype types and RSA traits in this natural population was analyzed, cultivars with Hap1 (n = 17) demonstrated heavier shoot and dry root weights, more lateral root numbers, and longer root lengths regardless of PRL and LRL. However, cultivars with Hap4 (n = 8) demonstrated the worst RSA traits for P absorption (Fig.s 4b-d).

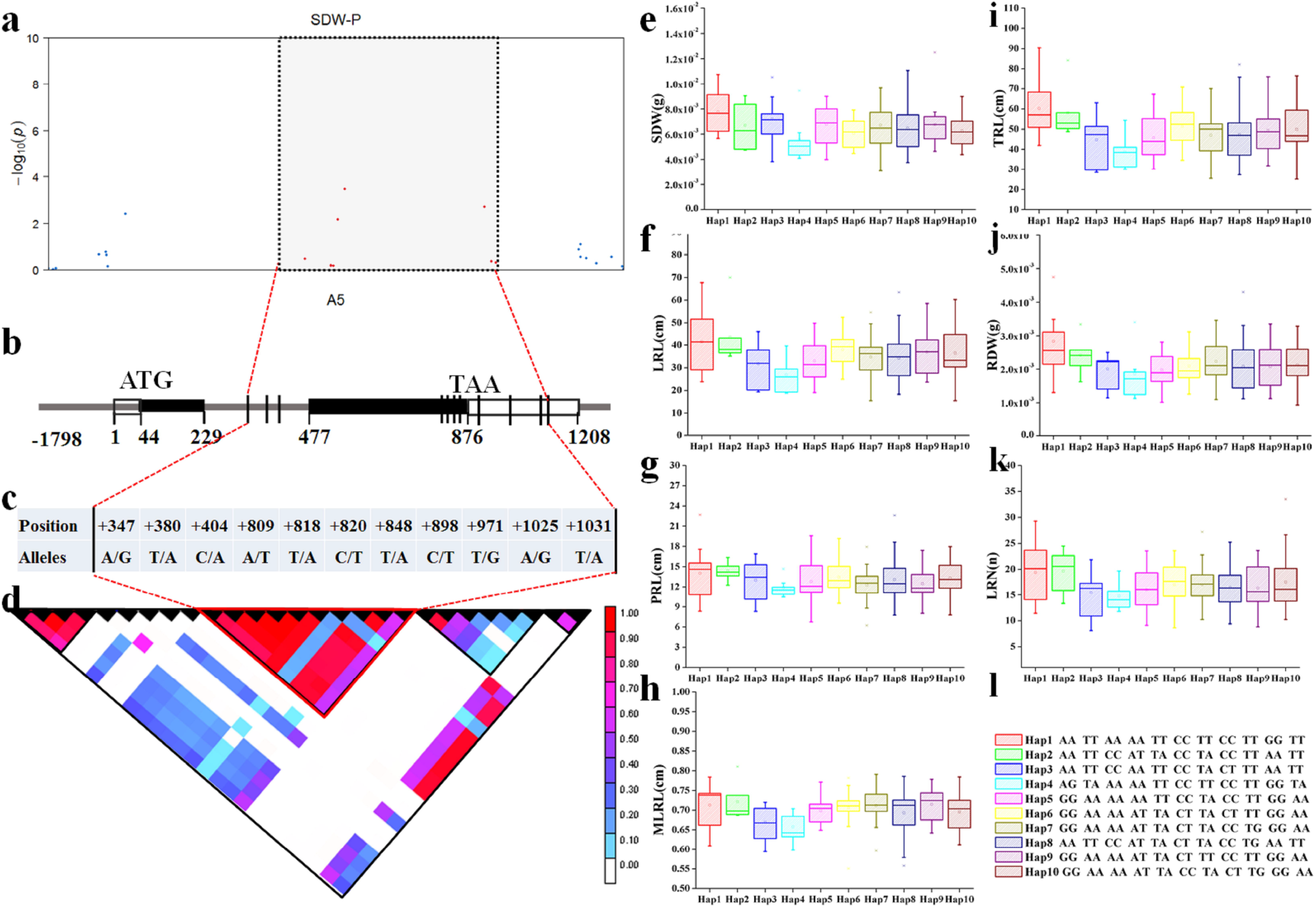

### BnGRP1 belongs to the glycine-rich protein family and responds to low P stress

To evaluate whether the Hap1 and Hap4 variation altered the P efficient and inefficiency function, the differences in sequence and relative expression between P efficient (with *BnGRP1*^*Hap1*^ type) and inefficiency (with *BnGRP1*^*Ha4*^ type) rapeseed cultivars were analyzed (Fig. S1). The *BnGRP1* on A5 revealed high expression in P efficient rapeseed ZS11 compared with P inefficient W10 in root tissues under low P stress. However, there were no differential expressions of homologous copies of *BnGRP1* on C5 between P efficient and inefficient cultivars in normal P conditions (Fig. S1). The *BnGRP1* encoded a protein of 291 amino acids, of which contained 181 glycines, i.e., a proportion of 62.2% (Fig. S2).

### BnGRP1^Hap1^ regulated low P tolerance in rapeseed

Previous studies have found *BnGRP1* to have a crucial role in environmental stress; however, low P stress was rarely reported. To assess whether the P efficient haplotype types *BnGRP1* ^*Hap1*^ contributed to low phosphorus tolerance and altered rapeseed function, this study conducted overexpression and CRISPR/Cas9-derived *BnGRP1*^*Hap1*^ knockout techniques on rapeseed. After investigating the *BnGRP1*^*Hap1*^ over-expression rapeseed lines *BnGRP1*-OE (Fig.s 5a, d, and e) at low P stress, the overexpression lines demonstrated longer root lengths, bigger surface areas, heavier root and shoot dry weights, and phosphorus content in root than control lines ‘Westar 10’ (Fig. 5b). During further analysis of the CRISPR/Cas9-derived *BnGRP1*^*Hap1*^ knockout mutations rapeseed *Bngrp1* under low P conditions, the mutations revealed shorter root lengths, lighter root, and shoot dry weight than the control line (Fig.s 5c, d, and e).

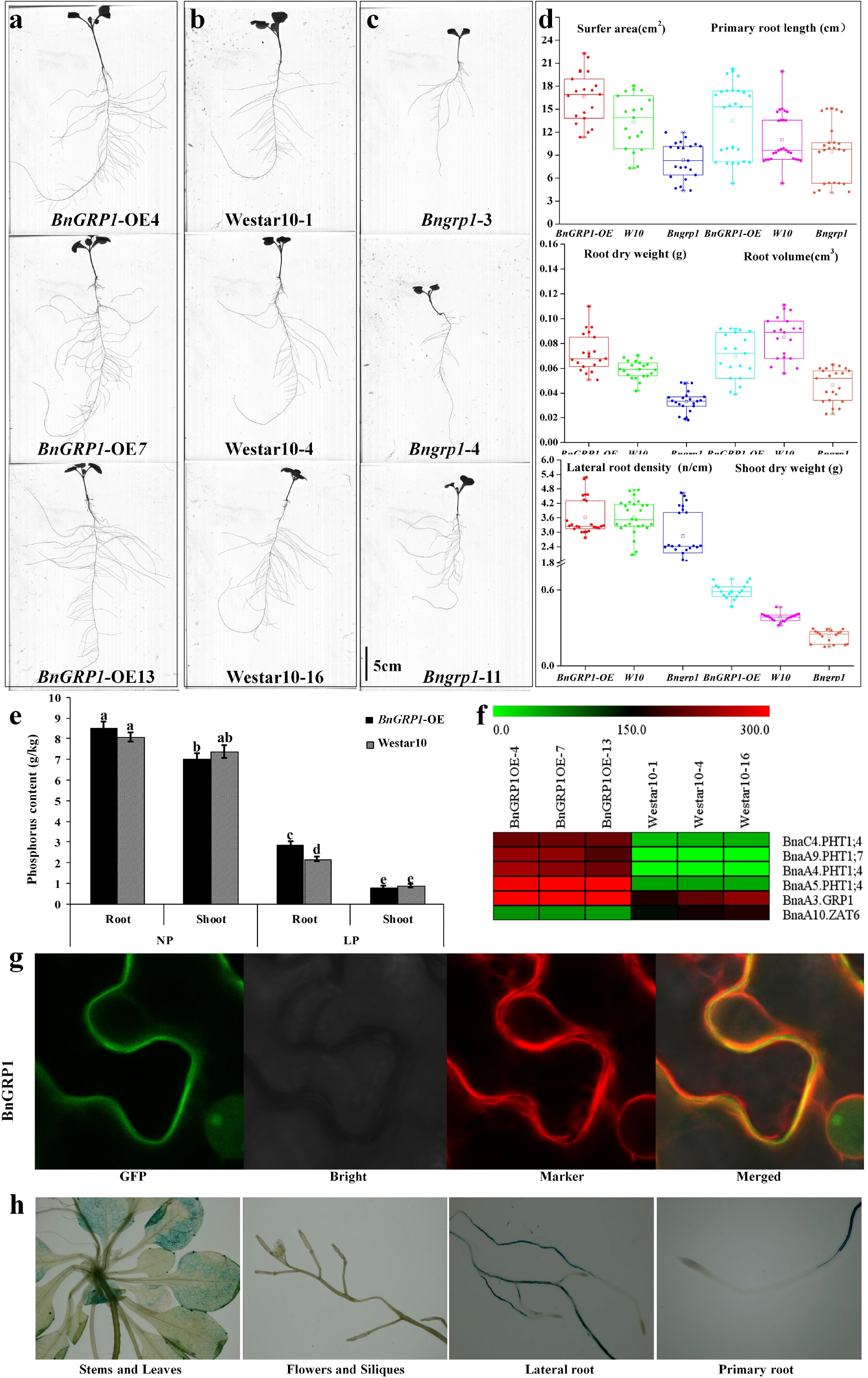

### BnGRP1 localization in cell wall and its specific expression in roots

Despite the numerous results confirming the contribution of *BnGRP1* to low P tolerance in rapeseed, the specific expression and subcellular localization were unclear. Therefore, GFP fluorescence was investigated in the cell wall after injecting the agrobacterium with recombinant *BnGRP1* vectors with cell membrane markers to the leaf bottom of *Nicotiana sanderae* (Fig. 5g). Moreover, there was a signal peptide site in the *BnGRP1* gene sequence, which could guide the transfer of newly synthesized proteins to the secretory pathway (Fig. S3). After investigating the *BnGRP1* GUS signal in *Arabidopsis* (35 days after germination), the tissue-specific expression was shown in the primary root and lateral root vascular system (without root tips), as well as leaf veins (Fig. 5h).

### BnGRP1 positively regulated the P uptake signaling pathway

Several transcription factors and functional genes are involved in the P uptake signaling pathways, such as *PHT1, PHR1, SPX1*, and *ZAT6*. However, no new genes have been reported in recent years. In this study, *BnGRP1* was identified to be related to RSA traits’ response to low P stress by GWAS. The over-expression phenotype and knockout mutation lines evidenced their role in P signaling. Here, rapeseed expression after *BnGRP1* over-expression using transcriptome sequencing revealed that with the increased expression of *BnGRP1*, the expression of *BnZAT6* was inhibited, but *BnPHT1* gene family members (*BnPHT1;4* and *BnPHT1;7*) were increased (Fig.s 5f and 6). The new model of *BnGRP1* was constructed to mediate that *PHT1* positively regulated the P uptake signaling pathway and contributed to low P tolerance.

Based on these findings, the genetic variation in *BnGRP1* might not only play a role in the constructed root system architecture but also be involved in the P uptake signaling pathway in *B. napus*.

## Discussion

As an important organ for nutrient absorption, the morphological change in RSA traits is an important indicator of P stress resistance (Lynch et al., 2011; Xu et al., 2022). The more RSA traits we identified, the more significant SNPs controlling low P tolerance were detected. In this study, 10 RSA traits were used to assess the response of rapeseed to low P stress. Based on the advantages of phosphorus-free paper culture (Wang et al., 2017), more RSA traits were identified, particularly regarding the root angle. In general, phosphorus is mainly distributed on the soil surface (Yamaji et al., 2017). During the evolution of rapeseed cultivation, rapeseed has developed a series of excellent root angles that benefit the absorption of surface P (Duan et al., 2021). However, due to difficulties in evaluating root angles in plants, knowledge about the genetic variations of root angles is rare. In total, 51 significant SNPs were identified as related to root angle under low P stress with controlling population structure in this study (Table S4).

In plants, low P tolerance-related traits are complex traits controlled by multiple genes (Wang et al., 2017). Several genes that respond to low P stress have not been identified in rapeseed. With the aid of *Brassica* 60K Illumina SNP array and re-sequencing, the genetic variations related to RSA traits under low P stress were detected in a natural population with 400 accessions. While previous studies only detected the genetic variations in the whole genome, the current study used GWAS and single chromosome associated study to ensure the accuracy of results (Fig. 3). Compared with GWAS, the single chromosome association study detected the LRL trait-related peak SNP (Tables S4 and S5). At the same time, the confidence intervals for the major candidate gene controlling RSA traits were further narrowed by facing the LD decay blocks (LD decay of A5 chromosome) of the significant SNP-related traits cluster. The QTLs related to different traits were mapped to the same significant SNPs, a finding attributable to pleiotropic influence (Jeon et al., 2021). Subsequently, transcriptome sequencing was used to analyze candidate genes that were differentially expressed in response to low phosphorus stress. This method was more efficient in identifying candidate genes than previous studies.

This study detected ten haplotype types based on 11 SNP variations on *BnGRP1* sequence using re-sequencing. The core region of the *BnGRP1* sequence was identified, as shown in Fig. 4. Based on the association between haplotype types and the phenotype of cultivars in the population, the P efficient and inefficient cultivars were defined. On the one hand, background rapeseed cultivars (Zhongshuang11 and Westar10) were used for over-expression and CRISPR/Cas9-derived knockout mutations. On the other hand, the cultivar with the Hap1 type enabled phosphorus efficiency in rapeseed breeding.

As a rich repetitive glycine gene family member, *BnGRP1* plays a critical role in plant stress resistance. GRP family genes are involved in cold and high salinity stress resistance through the osmotic regulation of cells mediated by the regulation of membrane oxidation (Tan et al., 2014; Znój et al., 2017). However, the role of *BnGRP1* in low P stress remained unclear. To gain insight into the contribution of *BnGRP1* towards low phosphorus tolerance in rapeseed, CRISPR/Cas9 overexpression was conducted in P efficient and P inefficient rapeseed cultivars, respectively. Similar to the function of *NtGRP3* promoting root length in *Nicotiana benthamiana* (Znój et al., 2017), *BnGRP1* increased root length in rapeseed in this study. The homologous gene in *Arabidopsis, AtGRP3*, is further evidence of its function in regulating root elongation and aluminum response pathways (Mangeon et al., 2016). In this study, two homologous copies of *BnGRP1* were detected, i.e., A5 and C5 copies. Although the functional copy could be identified by the significant association of SNPs, transcriptomes and qRT-PCR were used to improve the accuracy of A5 copy. The results showed that the *BnGRP1* on the A5 chromosome was only differentially expressed in root tissue under low P conditions (Fig. S1). Comparative sequencing analysis revealed a signal peptide recognition site on *BnGRP1* gene sequence (Fig. S3), indicating that the newly synthesized BnGRP1 was transferred through the secretory pathway.

Previous studies have mentioned the key role of phosphate transporter 1 (*PHT1*) genes in P uptake, transport, and reuse in plants (DiTusa et al., 2016; Dai et al., 2022. The expression of *PHT1* family is regulated by low P signaling and root tissue-specific expression. Similar to the expression pattern of *PHT1*, the *BnGRP1* expression was specifically regulated by low phosphorus in rapeseed root tissue (Fig. 5h). In *Arabidopsis, AtPHT1;4* was mainly expressed in root epidermis, cortex, pericycle, vascular cylinder, root cap, and lateral root under low P stress (Shin et al., 2004). Furthermore, P absorption capacity of *Atpht1;4* mutations was reduced by approximately 40% (Misson et al., 2004). After transferring the recombinant *BnGRP1* vector into *Arabidopsis*, GUS signals were invested in the vascular cylinder of the primary root, lateral root, and leaves vein in this study (Fig. 5h). After analysis of the over-expression of *BnGRP1*, the expression of *BnZAT6* was inhibited, but *BnPHT1* family members (*BnPHT1;4* and *BnPHT1;7*) were induced under low P stress (Fig.s 5f and 6). Overall, the findings of this study confirmed that *BnGRP1* mediates the up-regular expression of *PHT1* to enhance the rapeseed response to low P tolerance.

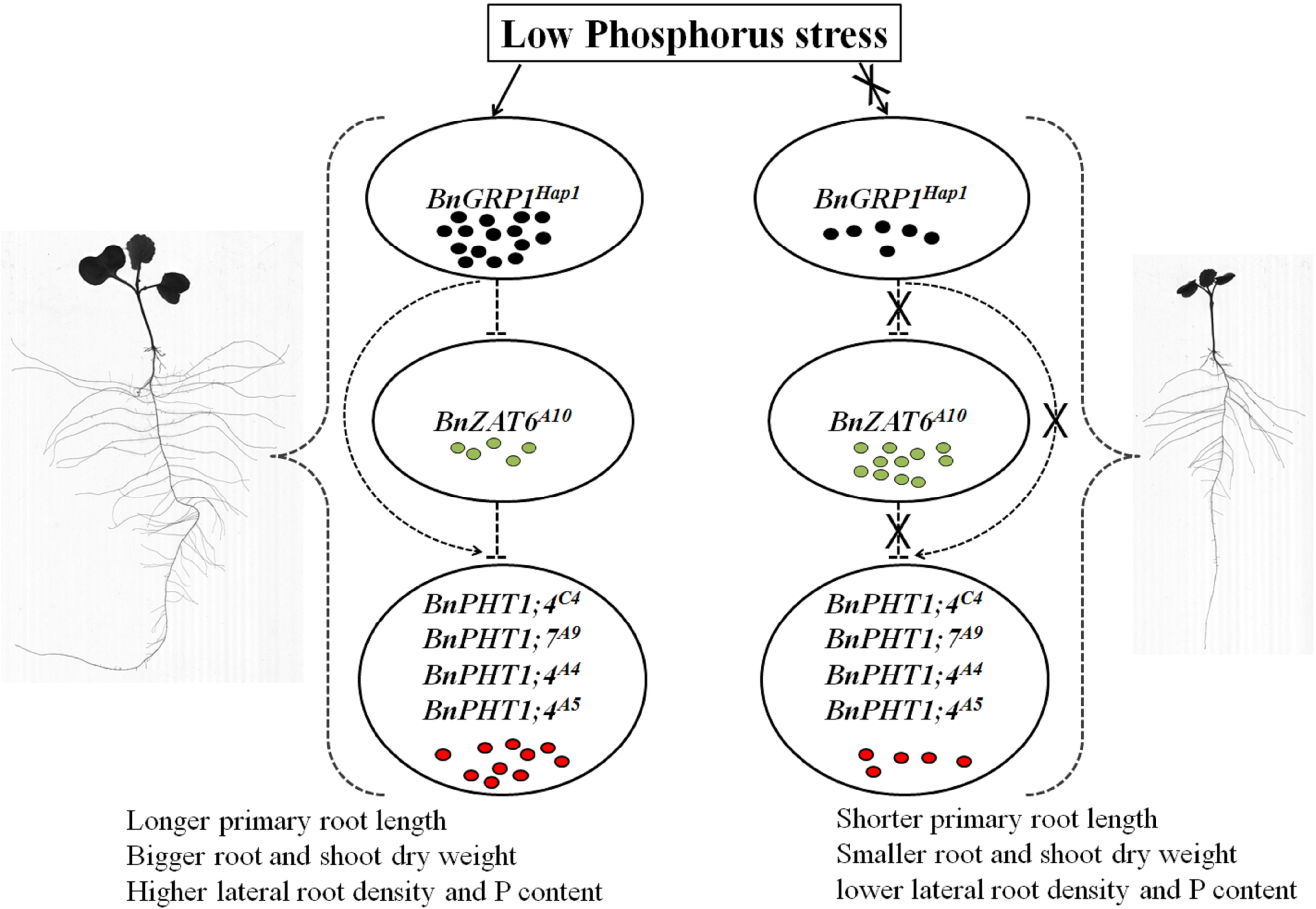

## Acknowledgments

This research was supported by the National Natural Science Foundation of China (32001575).

## Conflict of interest

The authors declare that they have no conflict of interest.

## Author contributions

Conceptual and experiment designs by P. X., H.L. and X.W.; Experiments were conducted by X.W., P.X. and K.X.; Data analysis performed by X.W., P. X., X. C., and Z. L; Reagents/materials/analysis tools were contributed by P.X., Z. L., and X.C. and the report was written by X.W., P.X. and L.Z. All the authors have commented, read and approved the final manuscript.

## Supporting information

Additional supporting information may be found online in the Supporting information section at end of the article.

**Figure S1**. The gene expression of BnGRP1 and their homologous copies in P efficient and inefficiency rapeseed cultivars. a. G1, *BnGRP1* on A5; b. G2, the homologous copies of *BnGRP1* on C5; ZS11, P efficient cultivar-Zhongshuang 11; W10, P inefficient cultivar-Westar 10; Root, root tissue; Shoot, young leaf and stem tissues. a, ab, b, c, cd was significant level at *p* < 0.05.

**Figure S2 Sequence analysis of *BnGRP1* protein in rapeseed and its homologous proteins in relative species**. a. the signal peptide sites of GRP1 sequence in rapeseed and relative species. b. the GRP1 protein sequence of rapeseed and relative species. Abbreviations: *Arabidopsis thaliana*-*At*; *Brassica rape*-*Br*; *Brassica cretica*-Bc; *Brassica oleracea-Bo*; *Raphanus sativus-RsG*; *Sinapis alba-Sa* and *Tarenaya hassleriana-Th*.

**Figure S3** The signal peptide site of *BnGRP1* gene sequence.

**Table S1**. The genotype of natural population using *Brassica* 60K Illumina Infinium SNP array.

**Table S2**. The population structure and genetic distance of natural population in this study.

**Table S3**. SNP number and LD decay of rapeseed association panel.

**Table S4**. The significant SNPs related to RSA traits in rapeseed natural population using genome-wide association study on 19 chromosomes.

**Table S5**. The genotype of natural population using Brassica 60K Illumina Infinium SNP array on A5 chromosome.

**Table S6**. The significant SNPs related to RSA traits in rapeseed natural population using genome-wide association study on A5 chromosome.

**Table S7**. The genes expression of LD confidence interval of peak SNP (SNP127-A5) on A5 chromosome.

**Table S8**. The SNP of *BnGRP1* sequence in natural population using re-sequencing.

**Table S9**. The primer sequence of CRISPR/Cas9 vector and plant transformation.

**Table S10**. The primer sequence of GUS vector and plant transformation.

## Notes

### Competing Interest Statement

The authors have declared no competing interest.

